# Residual errors in visuomotor adaptation persist despite extended motor preparation periods

**DOI:** 10.1101/2021.06.28.450124

**Authors:** Matthew Weightman, John-Stuart Brittain, R. Chris Miall, Ned Jenkinson

**Affiliations:** School of Sport, Exercise and Rehabilitation Sciences, University of Birmingham, Edgbaston, Birmingham, B15 2TT, UK; School of Psychology, University of Birmingham, Edgbaston, Birmingham, B15 2TT, UK; MRC-Versus Arthritis Centre for Musculoskeletal Ageing Research, University of Birmingham, Edgbaston, Birmingham, B15 2TT, UK; Centre for Human Brain Health, University of Birmingham, Edgbaston, Birmingham, B15 2TT, UK

**Author notes:** Corresponding Author: Matthew Weightman, School of Sport, Exercise and Rehabilitation Sciences, The University of Birmingham, Edgbaston, Birmingham, B15 2TT.

**Keywords:** Sensorimotor Adaptation, Motor Preparation, Mental Rotation

## Abstract

A consistent finding in sensorimotor adaptation is a persistent undershoot of full compensation, such that performance asymptotes with residual errors greater than seen at baseline. This behaviour has been attributed to limiting factors within the implicit adaptation system, which reaches a sub-optimal equilibrium between trial-by-trial learning and forgetting. However, recent research has suggested that allowing longer motor planning periods prior to movement eliminates these residual errors. The additional planning time allows required cognitive processes to be completed before movement onset, thus increasing accuracy. Here we looked to extend these findings by investigating the relationship between increased motor preparation time and the size of imposed visuomotor rotation (30°, 45° or 60°), with regards to the final asymptotic level of adaptation. We found that restricting preparation time to 0.35 seconds impaired adaptation for moderate and larger rotations, resulting in larger residual errors compared to groups with additional preparation time. However, we found that even extended preparation time failed to eliminate persistent errors, regardless of magnitude of cursor rotation. Thus, the asymptote of adaptation was significantly less than the degree of imposed rotation, for all experimental groups. Additionally, there was a positive relationship between asymptotic error and implicit retention. These data suggest that a prolonged motor preparation period is insufficient to reliably achieve complete adaptation and therefore our results provide support for the proposal that limitations within the implicit learning system contributes to asymptotic adaptation levels.

**New & Noteworthy:** Residual errors in sensorimotor adaptation are commonly attributed to an equilibrium between trial-by-trial learning and forgetting. Recent research suggested that allowing sufficient time for mental rotation eliminates these errors. In a number of experimental conditions, we show that while restricted motor preparation time does limit adaptation - consistent with mental rotation - extending preparation time fails to eliminate the residual errors in motor adaptation.

## Introduction

Sensorimotor adaptation has been extensively studied using visuomotor rotations (Cunningham, 1989; Krakauer, Ghilardi, & Ghez, 1999; Krakauer, Pine, Ghilardi, & Ghez, 2000). In this task, individuals adapt reaching movements to counter visual feedback that is rotated, for example, 30° or 45° from the hand’s position. However, adaptation is typically incomplete, and performance plateaus with errors a few degrees greater than at baseline, regardless of the rotation magnitude imposed (Fernandez-Ruiz, Wong, Armstrong, & Flanagan, 2011; Huberdeau, Haith, & Krakauer, 2015; van der Kooij, Brenner, van Beers, & Smeets, 2015; Vaswani et al., 2015; Weightman, Brittain, Punt, Miall, & Jenkinson, 2020; Langsdorf, Maresch, Hegele, McDougle, & Schween, 2021). State-space models of learning accurately capture this incomplete compensation (Smith, Ghazizadeh, & Shadmehr, 2006; Kording, Tenenbaum, & Shadmehr, 2007). In these models, error-driven learning and forgetting (or a reversion to baseline) work in opposition, and equilibrate below optimal performance, resulting in the commonly observed persistent error.

However, Vaswani and colleagues (2015) demonstrated that under certain conditions, individuals disengage from this limited error-dependent learning to attain greater task success. In one of their experimental conditions, Vaswani et al. (2015) ‘clamped’ visual feedback after an initial learning block, such that a small, fixed, visual error was presented, regardless of movement accuracy. They found that under these conditions, participants appeared to select an alternative, more exploratory learning policy, which enabled them to close the errors and even to overcompensate for the rotation (which differed from the partial reversion to baseline predicted by the state-space model). The authors suggested that the altered feedback distribution triggered alternative learning processes (and enabled the elimination of residual errors), which are usually suppressed during naturalistic feedback conditions.

More recently, Langsdorf et al. (2021) presented an alternative theory to explain why under normal, non-clamped environments, the central nervous system fails to eliminate residual errors during motor adaptation. They suggested an intrinsic speed-accuracy trade-off: where time consuming planning processes are interrupted by the imperative onset of movement, resulting in fast, but inaccurate movements. In their study, they showed that when an obligatory 2.5 second wait period was introduced between target presentation and movement initiation, participants were able to fully adapt to a 45° rotation, leaving no persisting errors at the end of learning. However, if this wait period was not enforced or was introduced at the end of the movement, when no planning was assumed to be taking place, participants failed to fully counteract the rotation. The authors highlighted mental rotation as a time-consuming cognitive process potentially involved in the planning of visuomotor adaptation.

For some time now, visuomotor adaptation has been framed as a combination of distinct learning processes (Mazzoni & Krakauer, 2006; Taylor, Krakauer, & Ivry, 2014; Bond & Taylor, 2015; McDougle, Bond, & Taylor, 2015; Voets, Panouilleres, & Jenkinson, 2020). These processes are customarily described as implicit and explicit. The implicit component adapts slowly and is driven by sensory prediction errors (Smith et al., 2006; Tseng, Diedrichsen, Krakauer, Shadmehr, & Bastian, 2007). The explicit component is a fast adaptation process and involves, among other processes, strategic re-aiming to counter the detected perturbation (Taylor & Ivry, 2011; Heuer & Hegele, 2008; McDougle et al., 2015). Such strategies are thought to include a form of metal rotation (McDougle & Taylor, 2019). Langsdorf et al. (2021) suggest that under naturalistic feedback conditions, parametric mental rotations from the target to the intended movement goal are prematurely terminated (despite no time constraints) and result in aimed movement trajectories falling short of the imposed rotation.

Behavioural and neurophysiological research suggests a role of mental rotation in the planning of movements aimed at angles away from visually defined targets (Georgopoulos & Massey, 1987; Bhat & Sanes, 1998; Georgopoulos, Lurito, Petrides, Schwartz, & Massey, 1989; Lurito, Georgakopoulos, & Georgopoulos, 1991; Pellizzer & Georgopoulos, 1993). Neuronal population vectors recorded in the monkey motor cortex gradually rotate from a stimulus direction to a cued movement direction during the planning of a reach (Georgopoulos et al., 1989; Lurito et al., 1991). Additionally, the completion of mental rotation tasks require long reaction times with larger magnitudes of rotation (Shepard & Metzler, 1971; Georgopoulos & Massey, 1987); a signature of cognitive strategies (Fernandez-Ruiz et al., 2011; Haith, Huberdeau, & Krakauer, 2015; McDougle & Taylor, 2019).

Given this knowledge, we aimed to replicate and extend the findings of Langsdorf et al. (2021), in order to further understand the roles of extended planning periods and mental rotation in attaining full adaptation. We designed an online visuomotor adaptation task, where participants had either long (2.5 seconds), medium (1 second) or short (0.35 seconds) enforced preparation periods between target presentation and movement onset, and were required to adapt to either a small (30°), moderate (45°) or large (60°) visuomotor rotation. Previous studies (Langsdorf et al., 2021) suggest that the 45° rotation would be fully corrected with the longest preparation period; errors for the middle interval would be greater; and a substantial residual error should be found for the shortest preparation interval. This negative relationship between preparation interval and residual error should be most obvious for the 60° rotation; the smallest rotation (30°) might allow full compensation even at the shortest preparation interval. In other words, the response time could be considered the sum of a mental rotation period linearly related to the rotation magnitude, and a fixed movement initiation period; if the sum exceeds the prescribed preparation interval, then residual errors should be seen.

## Materials and Methods

### Participants

A total of 200 participants were recruited via posters, study averts and through personal contact (aged 18-37 years, mean ± SD = 22.4 ± 3.5 years, 89 males); all gave informed consent before participating and either received no remuneration or received course credit which counted towards their university degree mark. The study was approved by the University of Birmingham ethics board (Science, Technology, Engineering and Mathematics Ethical Review Committee). Participants were self-reported as right-handed (181) or left-handed (19) and used their preferred hand to complete the task (all but 5 of the left-handed participants completed the task using their right hand). All participants had normal, or corrected to normal, vision and reported no history of neurological disease. Initially, 180 participants were pseudorandomised into one of nine experimental groups (each n = 20), which differed in the amount of preparation time provided (0.35, 1 or 2.5 seconds) and the magnitude of cursor rotation (30°, 45° or 60°). These participants formed the online arm of the study (Table 1). To avoid any possible confounds relating to online data collection and only after COVID-19 restrictions relating to in-person human testing were lifted, we also collected data from an additional experimental group who completed the task in a laboratory setting. Participants in this group experienced a 45° cursor rotation, with 2.5 seconds of preparation time (Table 1) - a closer replication of Langsdorf et al. (2021).

**Table 1:** Summary of experimental groups.

|  | 0.35 seconds |  |  | 1 second |  |  | 2.5 seconds |  |  |  |
| --- | --- | --- | --- | --- | --- | --- | --- | --- | --- | --- |
|  | 30° | 45° | 60° | 30° | 45° | 60° | 30° | 45° | 45° (lab) | 60° |
| N<br>(n male) | 20<br>(9) | 20<br>(6) | 20<br>(8) | 20<br>(11) | 20<br>(12) | 20<br>(4) | 20<br>(10) | 20<br>(13) | 20<br>(7) | 20<br>(9) |
| Mean age<br>(years, ± SD) | 23.5<br>(± 2.0) | 25.4<br>(± 3.4) | 25.9<br>(± 3.4) | 21.0<br>(± 1.1) | 21.5<br>(± 3.7) | 22.6<br>(± 5.0) | 20.8<br>(± 1.3) | 21.0<br>(± 1.6) | 20.2<br>(± 1.8) | 22.7<br>(± 3.7) |

### Task

#### Online

We developed a visuomotor adaptation task using the behavioural science experiment platform: PsychoPy (Peirce et al., 2019), which was implemented online via Pavlovia (https://pavlovia.org). Participants were required to have access to a desktop or laptop computer with internet connection and a mouse or trackpad to navigate through and complete the task (117/180 participants used a trackpad; 63/180 participants used a mouse). Participants were sent a link to access the task and entered a unique code which determined which combination of preparation time and rotation angle they would encounter. Participants were asked to sit in a location of their choosing, so that they could easily see their computer screen and comfortably reach and manipulate the mouse or trackpad. They made centre-out movements with either their mouse or on their trackpad from an on-screen central starting position to targets which appeared on an invisible circular array surrounding it (Figure 1a). They were told to make one fast and straight ‘shooting’ movement, aiming to get the on-screen cursor as close as possible to the target. Prior to starting the task, all participants completed a brief tutorial, which included the task instructions and aims. At this point, participants were asked to contact a researcher if they were unclear about any part of the task and/or had any questions relating to the instructions.

**Figure 1:**
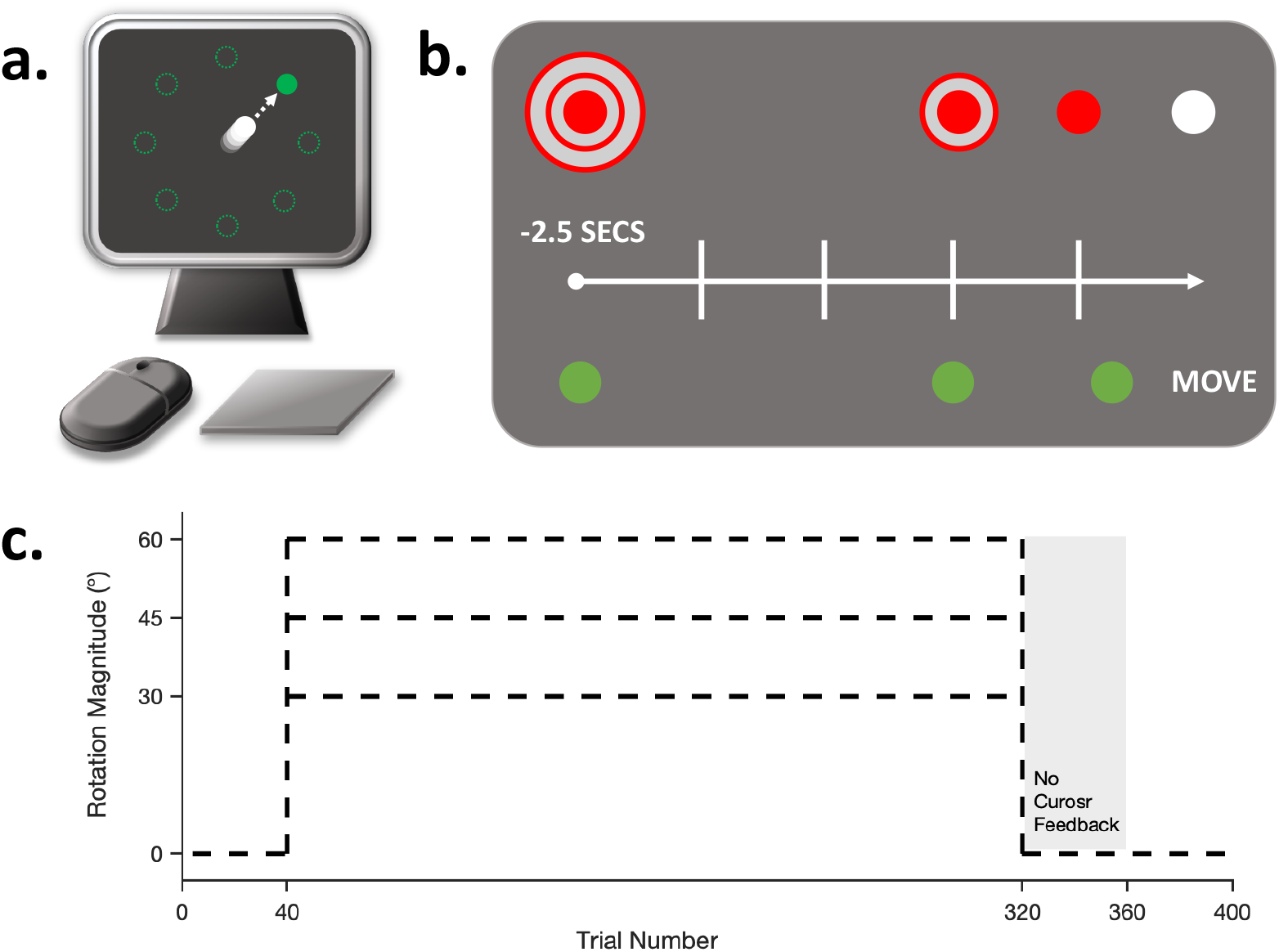
The experimental design. **a:** Participants used either a mouse or trackpad to direct an on-screen cursor towards targets presented radially around a central starting position. **b:** An example of the cursor countdown sequence and movement cue. At the beginning of each trial the (red) cursor would appear in the centre of the screen, flanked by two larger (red) rings (shown here on the top row). During the trial the two rings would disappear in sequence and the cursor would change colour to white. Participants were required to initiate their movement in synchrony with the cursor colour change, using the disappearing rings as a countdown. The target (shown on the bottom row) would appear either 2.5, 1 or 0.35 seconds prior to the movement cue, depending on group assignment. **c:** Time course of the study protocol. Participants completed 40 baseline trials, followed by 280 adaptation trials (where either a 30°, 45° or 60° cursor rotation was imposed), then 40 no-feedback trials and finally 40 de-adaptation trials.

#### Laboratory

Participants sat in an armless chair in front of a computer monitor (21.5 inch iMac, 60Hz refresh rate) and used a trackpad (Apple Magic Trackpad) to control the onscreen cursor. The trackpad measured 0.49-1.09 cm in height, 16 cm in width and 11.49 cm in depth and was fixed to the desk next to the computer screen. Crucially, the participants received the same instructions and completed the same tutorial before starting the task. The remaining task parameters were exactly as the online paradigm. All 20 participants were right-handed (self-report) and used their right hand to complete the task.

#### Protocol

At the beginning of every trial the cursor (red filled circle) would be located in the centre of the screen (grey background). The cursor was surrounded by two larger red annuli, which formed a bullseye-like formation (Figure 1b). At a set moment, the outer ring disappeared, followed by the inner ring 0.5 seconds later, and the cursor then changed colour from red to white after a further 0.5 seconds. Participants were told that this sequence of events should be treated as a countdown to movement and that they should aim to time their movement so that it was initiated in synchrony with the cursor changing colour from red to white (Figure 1b). Depending on the experimental group, the target (green filled circle) would appear either 2.5 seconds, 1 second or 0.35 seconds before the movement cue - thus creating three distinct preparation periods. The target could appear at any one of eight possible locations, equally spaced (separated by 45°) around an invisible circular array centred on the start position, and pseudorandomised so that the target appeared at each location once every cycle of eight trials. The position at which the cursor passed the invisible perimeter was displayed for 0.5 seconds as static open white circle, providing feedback of movement accuracy. After an inter-trial interval of 0.5 seconds the cursor would reset to the central position ready for the next trial. Participants were encouraged to move their hand back to a comfortable starting position after each trial. All elements of the visual display were scaled according to the size of the task window and thus were the same relative size for all participants. Consequently, the physical movement size required varied between participants depending on their set up and was not recorded beyond input device. If participants attempted to move 0.15 seconds before or after the go cue, or if movement time exceeded 0.25 seconds, the message “Response too fast/slow” appeared on the screen and the trial was aborted. These measures were to encourage fast movements, initiated correctly with the task instructions.

### Experimental Design

Participants first performed eight practice trials (one at each target location). These trials were performed with veridical cursor feedback and were used to familiarise the participants with the timing of their version of the task. The main task directly followed the practice trials and included four phases: baseline, adaptation, no-feedback and de-adaptation (Figure 1c). During baseline trials (n = 40) participants received veridical cursor feedback, such that the on-screen cursor moved in accordance with participants’ movements. Adaptation trials immediately followed, in which the cursor feedback was rotated 30°, 45° or 60° clockwise (depending on the experimental group, see below) relative to participants’ movement. This block lasted 280 trials, selected to ensure learning approached its asymptotic limit, based on pilot data collected prior to the present study. While there is some indication that full asymptotic saturation of learning was not achieved in all conditions, regression analysis of the last 40 trials of adaptation indicated that there was no significant trend remaining (all p > 0.19, with all slopes between 0.041 and −0.039) and thus we use the term ‘asymptote’ to refer to the final state reached at the end of the 280 adaptation trials. Following adaptation trials, participants performed no-feedback trials (n = 40). During these trials, participants were told to stop using any strategies they might have employed to achieve the task objectives and try to aim directly for the target. The cursor was hidden at all times during the trial, apart from when located in the central position for the countdown cue. End-point error was not displayed during the no-feedback phase. Veridical cursor feedback and display of end-point error were then restored for the final de-adaptation trials (n = 40).

There were ten participant groups (Table 1), to cover all combinations of preparation time (long: 2.5 seconds, medium: 1 second, short: 0.35 seconds) and the magnitude of cursor rotation (large: 60°, moderate: 45°, small: 30°), with the 2.5 seconds/45° group repeated in a laboratory setting. The preparation time was held constant throughout the whole task; the rotation was applied only during the adaptation phase.

### Data Analysis

There were three main outcome variables: reach angle, response time and movement duration. Our primary dependent variable - reach angle - was the angular difference between the target location and participants’ movement at end-point (i.e., the difference in angle between the vector linking the starting position and the target marker, and the vector linking the starting position and the point at which participants’ movement crossed the target circle perimeter). Participants were excluded from analysis if they either failed to follow task instructions (i.e., ignoring the rotated cursor and not adjusting their aiming direction) or if they violated the timing limits of the task on four successive trials (indicating a lack of concentration or distraction). 17 individuals were excluded based on this criterion and were replaced with new participants in order to achieve the desired sample size. Trials were deemed outliers and removed if they fell 2.5 standard deviations outside the group average on each trial. A total of 1.23% of all trials were removed from further analysis. Data from each participant were then averaged into bins of 4 trials to be used for visual representation and statistical analysis. Response time was defined as the time period between target presentation and movement onset (i.e., cursor movement breaking an invisible line immediately surrounding the starting position) and movement duration was defined as the time period between movement onset and when movement crossed the target circle. Reach angle data was not recorded for trials where the response or movement time limits were violated.

#### Statistical Analysis

Asymptotic levels of adaptation were defined as the mean error over the last 40 trials of the adaptation phase and compared between groups in a two-way ANOVA (Rotation Magnitude = 3 levels (30°/45°/60°) x Preparation Time = 3 levels (0.35 seconds/1 second/2.5 seconds)), with any significant main effects or interactions followed up with Bonferrroni-corrected multiple comparisons. Asymptotic differences from the imposed rotation magnitude were compared using a Wilcoxon Signed Rank test, after data from some groups failed normality checks (Shapiro-Wilk test). Mean reach angles across all baseline trials and during the first 16 no-feedback trials for each participant were used for baseline and retention comparisons. Data from the laboratory group were processed and analysed separately from the online data as this group was not factored into the original study design. In-lab data was compared to its online counterpart using unpaired t-tests (two-tailed) and from the degree of imposed rotation using a Wilcoxon Signed Rank test. All statistical analyses were carried out in MATLAB (R2018b) and SPSS (IBM, version 27). All ANOVAs were run as general linear models. The threshold for statistical significance was set at p < 0.05 and we report W, F, T and p-values, as well as effect sizes from ANOVAs (partial eta squared (ηp^2^)).

## Results

### Baseline performance does not differ between groups

Differences in reach angle during the baseline phase were assessed in a two-way ANOVA (Rotation Magnitude x Preparation Time), which revealed no significant main effects for Rotation Magnitude (F (2, 171) = 0.071, p = 0.93, ηp^2^ = 0.001), Preparation Time (F (2, 171) = 0.3, p = 0.75, ηp^2^ = 0.003), and no significant Rotation Magnitude*Preparation Time interaction (F (4, 171) = 0.45, p = 0.77, ηp^2^ = 0.1). This result suggests that all groups performed similarly during baseline trials (Figure 2) and thus, group differences are unlikely to influence performance later in the task. We felt this analysis important, given the limited experimenter input associated with online research.

**Figure 2:**
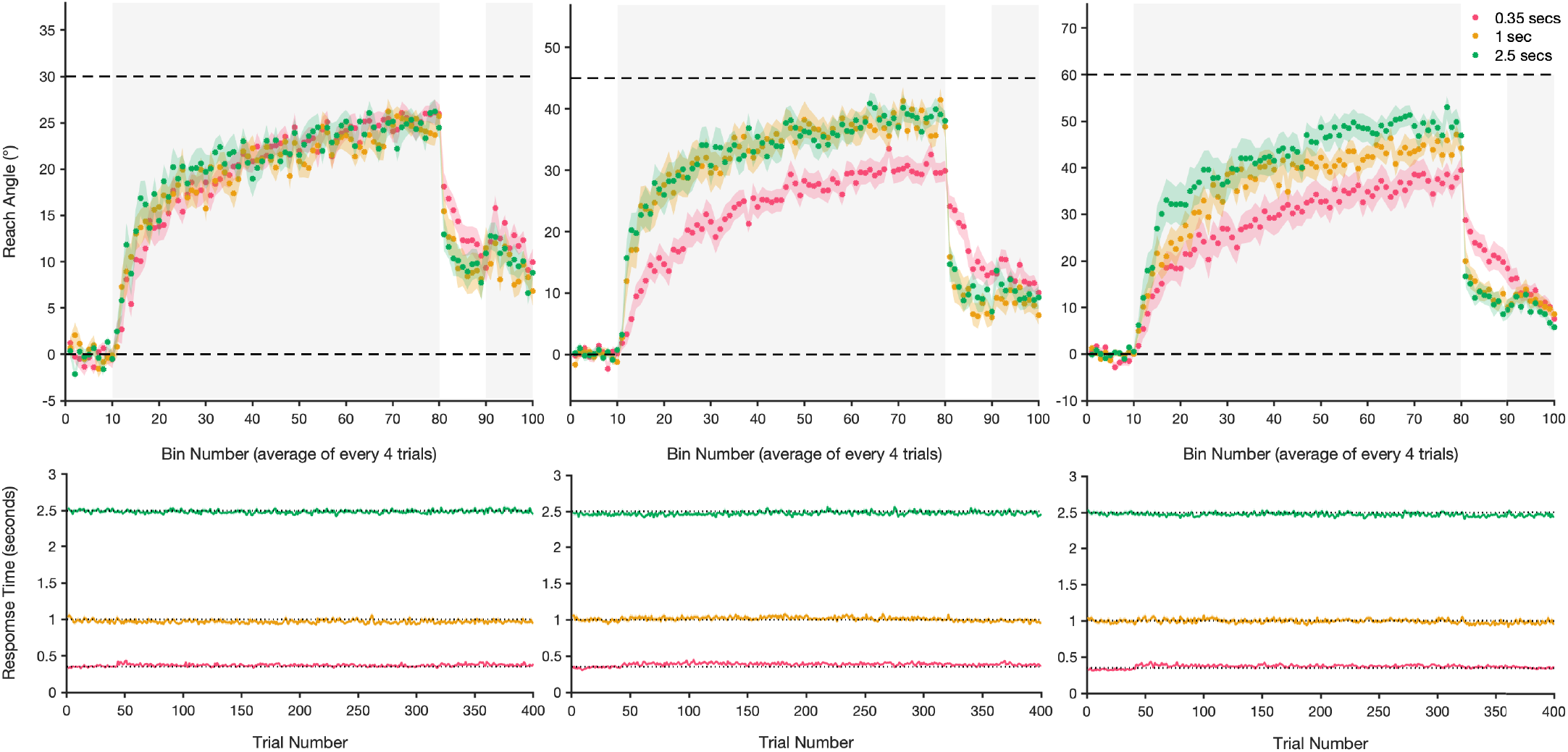
Mean reach angle and response time for each of the online experimental groups. Top panel: Reach error (± standard error, shaded region) is averaged every four trials into bins for (from left to right) the 30°, 45° and 60° groups, during baseline, adaptation (shaded grey background), no feedback and de-adaptation trials (shaded grey background). Zero degrees and the magnitude of imposed rotation are shown by horizontal dashed lines. Bottom panel: Mean response time (time between target presentation and movement initiation), ± standard error (shaded region) for each group. Note, preparation times were predetermined at 0.35, 1 or 2.5 seconds (dotted lines) and were tightly controlled.

### The effect of differing preparation time on asymptotic levels of adaptation

A two-way ANOVA comparing asymptotic levels of adaptation revealed significant main effects for Rotation Magnitude (F (2, 171) = 121.01, p < 0.001, ηp^2^ = 0.59), Preparation Time (F (2, 171) = 15.17, p < 0.001, ηp^2^ = 0.15) and a significant Rotation Magnitude*Preparation Time interaction (F (4, 171) = 5.24, p < 0.001, ηp^2^ = 0.11). Multiple comparisons revealed that in the 45° rotation group, there were some differences between learning asymptotes, with regards to preparation time provided. The 0.35 second preparation time group displayed impaired adaptation compared to the 1 and 2.5 second groups (both p < 0.001), with no further differences between the latter two conditions (p > 0.99). However, contrary to Langsdorf et al. (2021), all final levels of adaptation significantly differed from 45° (0.35 seconds: W = −210.0, p < 0.001, 95% CI = [−16.72, −11.32]; 1 second: W = −200.0, p < 0.001, 95% CI = [−9.71, −2.50]; 2.5 seconds: W = −210.0, p < 0.001, 95% CI = [−9.33, 3.18]). The 0.35 seconds preparation time group compensated 67.3% for the rotation, with the 1 and 2.5 second groups achieving 85.1% and 86.1% compensation respectively (Figures 2 & 3), suggesting that additional preparation time allowed for greater compensation but not the elimination of residual errors.

**Figure 3:**
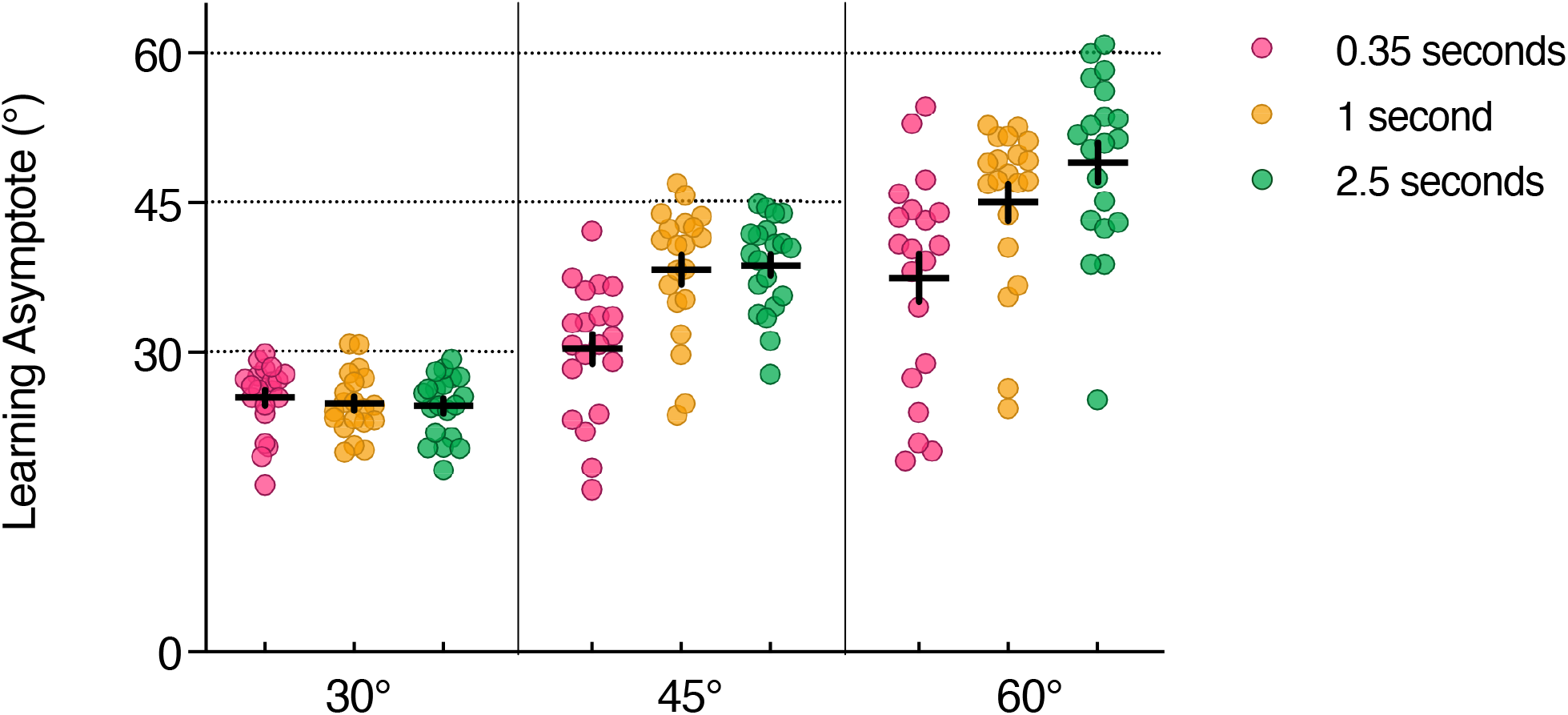
The asymptotic levels of adaptation for each participant in each of the rotation magnitude and preparation time groups (online experiment). Filled circles represent the adaptation asymptote for each participant (defined as the average of the last 40 adaptation trials), with mean values and standard error depicted by horizontal and vertical lines respectively. Dashed horizontal lines depict the imposed cursor rotation for each group.

Similarly, for the 60° rotation condition, shortening planning time to 0.35 seconds significantly reduced the final level of adaptation compared to both 1 and 2.5 second preparation periods (0.35 seconds vs 1 second: p = 0.001, 0.35 seconds vs 2.5 seconds: p < 0.001). However, there was no difference between the 1 and 2.5 second groups with respect to final adaptation levels (p = 0.17). Participants averaged 62.4% compensation with 0.35 seconds preparation time, 75.0% with 1 second preparation time and 81.7% with 2.5 seconds preparation time (Figures 2 & 3). Additionally, all asymptotic levels of adaptation significantly differed from the imposed 60° rotation (0.35 seconds: W = −210, p < 0.001, 95% CI = [−31.14, −16.02]; 1 second: W = −210.0, p < 0.001, 95% CI = [−16.27, −10.27]; 2.5 seconds: W = −206.0, p < 0.001, 95% CI = [−16.76, −6.44]).

In the 30° rotation group, there were no differences in asymptotic error between any of the different preparation time groups (all p > 0.99, Figure 2). Furthermore, all final levels of adaptation differed significantly from 30° (0.35 seconds: W = −210.0, p < 0.001, 95% CI = [−5.35, −2.39]; 1 second: W = −204.0, p < 0.001, 95% CI = [−6.87, −2.98]; 2.5 seconds: W = −210.0, p < 0.001, 95% CI = [−8.09, −3.25]), with adaptation reaching 84.7%, 82.8% and 82.0% of the optimum, respectively (Figure 3).

In summary, our results support previous work suggesting that mental rotation contributes an interval to the planning process which is proportional to rotation magnitude, and that short periods of preparation interrupt planning, leading the greater residual errors. However, for the smallest rotation, we found no relationship between preparation time and error, suggesting that mental rotation was completed within the allowed time. But even in these circumstances, there were significant residual errors. In fact, in opposition to results reported by Langsdorf et al. (2021), all groups displayed incomplete adaptation regardless of the preparation time provided.

### Restricting motor preparation time resulted in a stronger implicit retention

Implicit retention (mean reach angle in the first 16 no-feedback trials, two cycles of reaches to each target location) was compared between groups in a two-way ANOVA (Rotation Magnitude x Preparation Time). Results from the ANOVA revealed a significant main effect of Rotation Magnitude (F (2, 171) = 6.71, p = 0.002, ηp^2^ = 0.07) and Preparation Time (F (2, 171) = 18.33, p < 0.001, ηp^2^ = 0.18), but no significant interaction (F (4, 171) = 1.08, p = 0.37, ηp^2^ = 0.03). Multiple comparisons showed that participants in the restricted 0.35 second motor preparation time groups displayed greater retention than both the 1 second (p < 0.001) and 2.5 second (p < 0.001) groups, with no differences between the latter two (p > 0.99).

### Input device and handedness had no effect on late adaptation levels

In order to determine whether the input device used (trackpad or mouse), or the handedness of the participant had any effect on our main outcome variable, reach angle, we ran an additional mixed design ANOVA (Rotation Magnitude x Preparation Time) with factors for both input device and handedness (online groups only). The ANOVA revealed no significant main effect of Input Device (F (1, 150) = 0.9, p = 0.35, ηp^2^ = 0.006) or Handedness (F (1, 150) = 1.12, p = 0.29, ηp^2^ = 0.007). In fact, no interaction that included the term Input Device revealed significance (Rotation Magnitude*Input Device: F (2, 150) = 0.36, p = 0.7, ηp^2^ = 0.005; Preparation Time*Input Device: F (2, 150) = 0.23, p = 0.79, ηp^2^ = 0.003; Rotation Magnitude*Preparation Time*Input Device: F (4, 150) = 0.3, p = 0.88, ηp^2^ = 0.008; Supplementary Figure 1, see https://doi.org/10.6084/m9.figshare.14797926). Similarly, all interaction effects with Handedness were also non-significant (Rotation Magnitude*Handedness: F (2, 150) = 1.55 p = 0.22, ηp^2^ = 0.02; Preparation Time*Handedness: F (2, 150) = 0.75, p = 0.48, ηp^2^ = 0.01; Rotation Magnitude*Preparation Time*Handedness: F (2, 150) = 0.12, p = 0.89, ηp^2^ = 0.002). These analyses corroborate our belief that neither the handedness of the participants, nor the input devices they used, confound our results.

### There were no differences in task performance between the online and in-laboratory groups

To confirm the validity of our results collected using online methodology, we collected data from an additional group of participants who performed the experiment in a laboratory with 2.5 seconds of preparation time and a 45° cursor rotation. Data from this additional in-lab experimental group were compared the equivalent online group using unpaired t-tests (two-tailed) and revealed no significant differences in baseline, asymptotic or retention performance (t(38) = 0.89, p = 0.38; t(38) = 0.35, p = 0.73; t(38) = 1.78, p = 0.084, respectively). Participants in this group compensated 87.2% for the rotation and importantly, the final level of adaptation was also significantly different from 45° (W = −196.0, p < 0.001, 95% CI = [−8.35, −3.1]), replicating the effect found consistently across our online groups (Figure 4). We also found no differences in variance between the laboratory and online groups in each of the experimental phases (F-tests: F(19, 19) = 1.9, p = 0.17; F(19, 19) = 1.04, p = 0.93; F(19, 19) = 1.31, p = 0.56).

**Figure 4:**
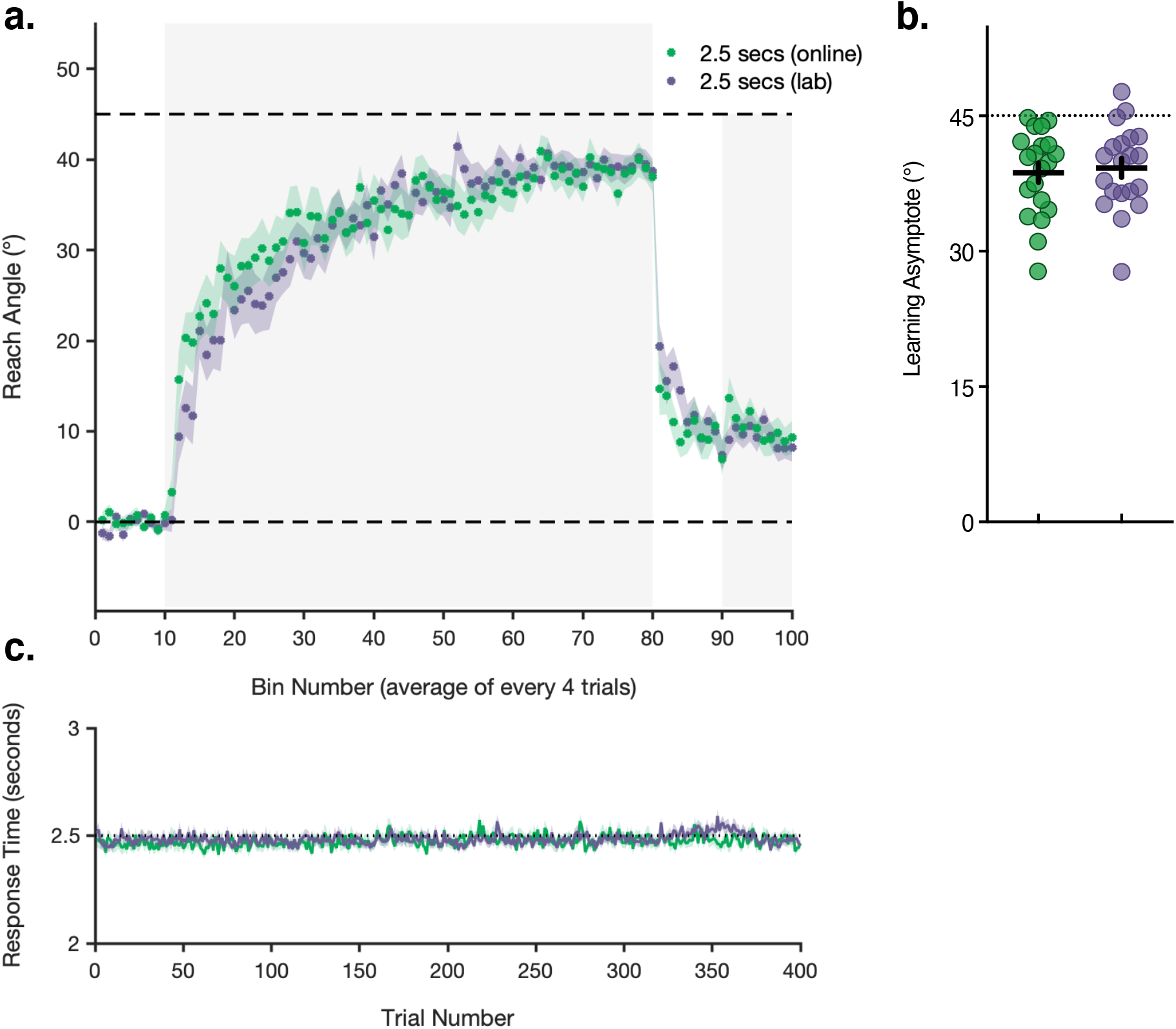
Mean reach angle, response time and learning asymptote for the in-laboratory experimental group. **a:** Reach error (± standard error, shaded region) is averaged every four trials into during baseline, adaptation (shaded grey background), no feedback and de-adaptation trials (shaded grey background) for each group. Zero degrees and the magnitude of imposed rotation are shown by horizontal dashed lines. **b:** The asymptotic level of adaptation for each participant. Filled circles represent the adaptation asymptote for each participant (defined as the average of the last 40 adaptation trials), with mean values and standard error depicted by horizontal and vertical lines respectively. Dashed horizontal lines depict the imposed cursor rotation. **c:** Mean response time (time between target presentation and movement initiation), ± standard error (shaded region). Preparation time was predetermined and tightly controlled at 2.5 seconds.

## Discussion

We aimed to test the assumption that increased motor preparation periods may allow for more complete adaptation during visuomotor rotation tasks. As such, we hypothesised that shorter preparation periods would be sufficient to fully counteract a 30° rotation, as a small rotation would require less (and therefore quicker) mental rotation before motor execution. We then predicted that this effect would scale with an increased cursor rotation, so that 45° and 60° would require greater mental rotation and thus more time to fully compensate. Indeed, we did find that restricting planning time for moderate and larger rotations resulted in impaired final adaptation levels and greater implicit retention compared to those groups with more preparation time. However, contrary to research from Langsdorf et al. (2021), we found no evidence to suggest that extended motor preparation periods allow for the elimination of residual errors observed during late stages of adaptation, as all groups displayed final adaptation performance significantly lower than the imposed cursor rotation.

Hence, we failed to replicate the results from Langsdorf et al. (2021), who found that enforcing a 2.5 second planning period between target presentation and reach onset allowed individuals to fully counteract a 45° visuomotor rotation, leaving no residual mean error at asymptote. The authors suggested that the extended planning period allowed for time-consuming cognitive processes, such as mental rotation, to be completed prior to the go-signal and resulted in complete compensation. This theory, centred around a speed-accuracy trade-off, seems plausible given that reaction times increase in a linear fashion when subjects are asked to reach at increasing angles from a target - a relationship that mimics the behaviour seen in mental rotation tasks (Shepard & Metzler, 1971; Georgopoulos & Massey, 1987; McDougle & Taylor, 2019). Accordingly, brain regions activated in both visuomotor adaptation and mental rotation tasks have also been shown to overlap (Anguera, Reuter-Lorenz, Willingham, & Seidler, 2010), suggesting that mental rotation contributes to visuomotor adaptation. Recent evidence investigating preparatory activity in the motor cortex of non-human primates further suggests the necessity for motor planning time in visuomotor adaptation (Vyas, O’Shea, Ryu, & Shenoy, 2020). In addition, Langsdorf et al. (2021) found that delaying movement initiation resulted in an overcompensation of the imposed rotation, similar to the overshoot in reach estimation shown by Georgopoulos and Massey (1987), suggesting similar cognitive rotation strategies may have been in play.

Notwithstanding these studies, we found that on average participants’ learning plateaued before eliminating error, with asymptotic learning levels statistically different from the degree of imposed rotation for all rotational groups (30°, 45°, 60°), regardless of the amount of preparation time enforced. These findings are consistent with previous reports which have suggested that sustained errors during visuomotor adaptation are a product of the implicit learning system; either suppressing alternative learning mechanisms which may be able to overcome persisting errors (Shmuelof et al., 2012; Vaswani et al., 2015) and/or modulating the system’s sensitivity to errors dependent on their size, variability and history (Albert et al., 2021). It should be noted however, that despite regression analysis suggesting the data had reached an asymptote, there is a possibility that with a longer adaptation session, or with multiple sessions, participants may have been able to reduce their error further and get closer to complete compensation. Yet, given the degree of error remaining at the end of these 280 trials, and the non-significant slopes of the regression analyses, it is unlikely that much greater adaptation would be achieved within the bounds of Langsdorf et al’s. (2012) study protocol (i.e., with an additional 40 trials). In support of this, estimated asymptotic levels of adaptation modelled from existing mean group data are in close correspondence with the values calculated in our original analysis and suggest all groups would asymptote below the degree of imposed rotation (Supplementary Figure 2; see https://doi.org/10.6084/m9.figshare.17136857).

One factor that may explain the discrepancy between our results and those of Langsdorf et al. (2021), is the level of experimental control. Our study was conducted online, with participants completing the task unsupervised at home, whereas Langsdorf et al. (2021) conducted a laboratory-based study. The disparities associated with these environments may have contributed to small but significant differences in motivation, attention and behaviour. That said, data from our additional in-laboratory group is at odds with this reasoning. We found no difference in adaptive performance at any phase of the task between the in-lab and corresponding online group, with participants in both groups failing to achieve full compensation. This finding supports previous literature which suggests that kinematic data and results from online motor learning studies are analogous with those collected in a controlled lab environment (Tsay, Lee, Ivry, & Avraham, 2021) and that online tasks perform well in terms of accuracy and precision with regards to timing of visual stimuli and response capture (Bridges, Pitiot, MacAskill, & Peirce, 2020). Furthermore, our data are typical of many previous lab-based visuomotor adaptation studies, with comparable levels of final performance (Huberdeau et al., 2015; Morehead, Qasim, Crossley, & Ivry, 2015; Vaswani et al., 2015; Neville & Cressman, 2018; Weightman et al., 2020; Weightman, Brittain, Miall, & Jenkinson, 2021). With all of these results in mind, it seems improbable that the failure to eliminate residual errors (and replicate results from Langsdorf et al., 2021) was a consequence of online data collection methods.

An additional difference between the two studies, which may have had some bearing on the results, is the nature of response cueing used. We used a visual countdown sequence which enabled participants to accurately synchronise their movements with the go-signal and ensured a tight coupling between response times and the preparation time groupings. In contrast, Langsdorf et al. (2021) displayed the target and instructed participants to wait until they heard a tone before responding. This difference in protocol resulted in response times closer to 3 seconds. One may argue that more than 2.5 seconds are required to fully prepare for a 45° rotated reach. This is unlikely however, as we show that even a 30° angle was not fully compensated, and there was no evidence for a relationship with the preparation interval for those groups. Previous research has also shown that mental rotation and aiming towards angles up to 90° can be achieved in ~1 second (McDougle & Taylor, 2019). There is perhaps an argument to suggest that attending to the countdown sequence may have interfered with mental rotation and other cognitive planning processes, by, for instance vying for attentional resources and thus contributing to a consistent reach undershoot (Taylor & Thoroughman, 2007; Galea, Sami, Albert, & Miall, 2010). Nevertheless, prior studies using auditory response cueing have not cited attention to the timed-response sequence as a potential confound (Haith et al., 2015; Leow, Gunn, Marinovic, & Carroll, 2017; McDougle & Taylor, 2019).

Despite participants failing to eliminate residual errors during late adaptation, our data do include some hallmarks of a speed-accuracy trade-off. We found that motor adaptation was impaired in the 45° and 60° rotational groups, during both early and later stages, when planning time was restricted to 0.35 seconds. This result is in line with previous studies who report reduced error compensation when preparation times are limited (Fernandez-Ruiz et al., 2011; Leow, Gunn, et al., 2017; Leow, Marinovic, Riek, & Carroll, 2017; Albert et al., 2021) and may reflect the suppression of explicit re-aiming/cognitive strategies. Additionally, impaired adaptation associated with restricted preparation times was coupled with an increased retention during no-feedback trials, which is consistent with the idea that learning involved more implicit, procedural processes (Fernandez-Ruiz et al., 2011; Haith et al., 2015). The greatest retention was seen in the 0.35 seconds / 60° group, which implies that adaptation in this group may have been weighted more so towards implicit processes – also highlighted by the larger residual errors at the end of learning. This suggestion is reinforced by the equal retention seen for the 2.5 second condition, across all three rotation magnitudes. In this case, the residual errors are approximately equal, because the preparation time exceeded that required for mental rotation. However, these assumptions should be treated with some caution after recent commentary on how the dichotomy of implicit and explicit components of motor adaptation are inferred (Hadjiosif & Krakauer, 2021).

Although not statistically evident, a speed-accuracy trade-off may also be apparent in the 60° rotational group (Figure 3). On average, there was a linear increase in learning asymptote as preparation time increased, which may reflect how mental rotation of the intended movement angle increases the accuracy of reaches (McDougle & Taylor, 2019; Langsdorf et al., 2021). Addition of further adaptation trials might exaggerate these asymptotic differences, until statistically significant, yet data from our other rotational groups and previous literature would suggest that errors would still persist. To achieve full compensation, it is possible that explicit information about the nature of rotation ought to be provided (McDougle & Taylor, 2019) or strategies primed using methods such as verbal aiming reports (Bond & Taylor, 2015; Brudner, Kethidi, Graeupner, Ivry, & Taylor, 2016; Wilterson & Taylor, 2019; Albert et al., 2021; Langsdorf et al., 2021), which are inherently coupled with increased reaction/plan times.

In summary, our data suggests that extending motor preparation and planning periods alone is insufficient to eliminate residual errors during visuomotor adaptation, irrespective of the size of imposed cursor rotation. While increased preparation time may help to improve error reduction at larger rotation magnitudes, our results suggest there remains a limit at which learning saturates at asymptote, perhaps only overcome with priming of explicit strategies, further instruction or changes to experimental variables. We also provide further support of the use of online data collection methods in the study of motor control and learning. Understanding why the central nervous system fails to fully adapt movements in response to environmental changes may be key when aiming to optimise rehabilitation protocols following brain injury or disease.

## Acknowledgements

This work was funded by the MRC-Versus Arthritis Centre for Musculoskeletal Ageing Research (CMAR). RCM was also partly funded by the Royal Society, Leverhulme and Wellcome Trust.

## Author Contributions

MW, JSB and NJ conceived and designed the study with input from RCM. MW collected the data. MW, JSB, RCM and NJ analysed and interpreted the data. MW wrote the first draft of the paper. JSB, RCM and NJ reviewed and edited the paper and all authors agreed with the final submitted version.

The authors declare no competing interests.

