## Supplementary Figure. 1 for "Residual errors in visuomotor adaptation persist despite extended motor preparation periods" (from another version of this preprint: version 1, 2021-06-28)

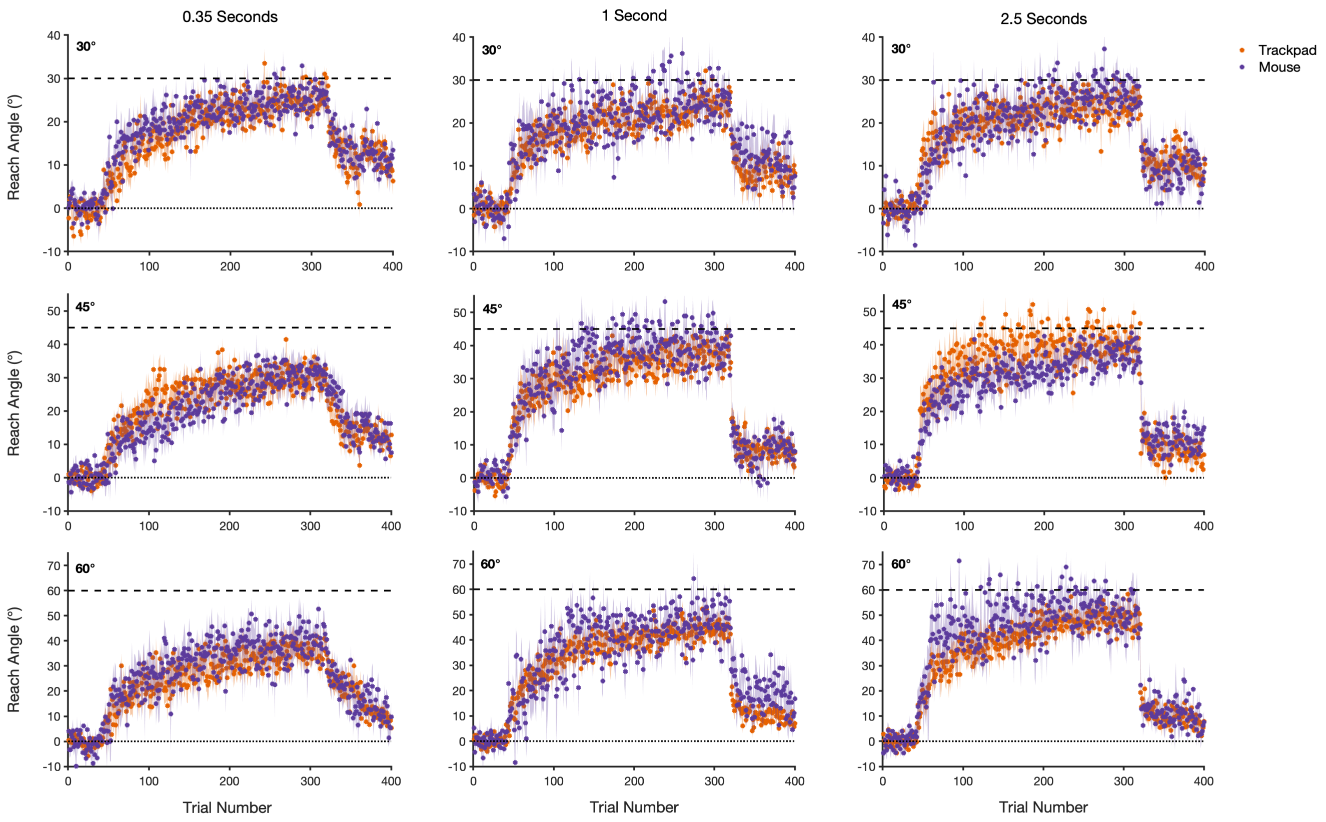


**Supplementary Figure. 1:** *Mean reach angle (coloured markers, ± standard error; shaded region) during baseline, adaptation, no-feedback and de-adaptation trials for all the experimental groups. Data is separated by the input device used to complete the task (mouse or trackpad). Zero degrees and the magnitude of imposed rotation shown as a dotted and dashed line respectively.*

*A 3x3x2 mixed-design ANOVA (Rotation Magnitude x Preparation Time x Input Device) revealed no significant main effect of Input Device ($F(1, 179) = 1.13$, $p = 0.29$, $\eta p^2 = 0.007$) and no Rotation Magnitude*Input Device ($F(2, 179) = 0.53$, $p = 0.59$, $\eta p^2 = 0.006$), Preparation Time*Input Device ($F(2, 179) = 0.53$, $p = 0.59$, $\eta p^2 = 0.006$) or three-way interaction ($F(4, 179) = 0.38$, $p = 0.82$, $\eta p^2 = 0.009$).*
